# Whole-genome sequencing of three native cattle breeds originating from the northernmost cattle farming regions

**DOI:** 10.1101/390369

**Authors:** Melak Weldenegodguad, Ruslan Popov, Kisun Pokharel, Innokentyi Ammosov, Ming Yao, Zoya Ivanova, Juha Kantanen

**Author notes:** Correspondence: Juha Kantanen.

## Abstract

Northern Fennoscandia and the Sakha Republic in the Russian Federation represent the northernmost regions on Earth where cattle farming has been traditionally practiced. In this study, we performed whole-genome resequencing to genetically characterize three rare native breeds Eastern Finncattle, Western Finncattle and Yakutian cattle adapted to these northern Eurasian regions. We examined the demographic history, genetic diversity and unfolded loci under natural or artificial selection. On average, we achieved 13.01-fold genome coverage after mapping the sequencing reads on the bovine reference genome (UMD 3.1) and detected a total of 17.45 million single nucleotide polymorphisms (SNPs) and 1.95 million insertions-deletions (indels). We observed that the ancestral species (*Bos primigenius*) of Eurasian taurine cattle experienced two notable prehistorical declines in effective population size associated with dramatic climate changes. The modern Yakutian cattle exhibited a higher level of within-population variation in terms of number of SNPs and nucleotide diversity than the contemporary European taurine breeds. This result is in contrast to the results of marker-based cattle breed diversity studies, indicating assortment bias in previous analyses. Our results suggest that the effective population size of the ancestral Asiatic taurine cattle may have been higher than that of the European cattle. Alternatively, our findings could indicate the hybrid origins of the Yakutian cattle ancestries and possibly the lack of intensive artificial selection. We identified a number of genomic regions under selection that may have contributed to the adaptation to the northern and subarctic environments, including genes involved in disease resistance, sensory perception, cold adaptation and growth. By characterizing the native breeds, we were able to obtain new information on cattle genomes and on the value of the adapted breeds for the conservation of cattle genetic resources.

## Introduction

During their 8,000-10,000 years of domestication, taurine cattle (*Bos taurus*) have adapted to a wide variety of biogeographic zones and sociocultural environments as a result of natural and human-derived selection (Felius, 1995). Fennoscandia along with northwestern Russia and the region of Sakha (Yakutia) in eastern Siberia, are the northernmost territories where cattle farming has had a relatively long tradition as the livelihood of local people (Kopoteva and Partanen, 2009; Bläuer and Kantanen, 2013; Cramp et al., 2014; Egorov et al., 2015). In prehistoric and historic times, animal husbandry faced several challenges in these northern climatic conditions, such as short summers and limited vegetation resources for feeding during the long winters, and this practice required well-adapted animals that were suited to the available environmental resources and socioeconomic and cultural conditions (Kantanen et al., 2009a; Bläuer and Kantanen, 2013; Egorov et al., 2015).

Cattle breeds such as Eastern Finncattle, Icelandic cattle, Swedish Mountain cattle, Yakutian cattle and other northern native cattle breeds are assumed to have their origins in the near-eastern domesticated taurine cattle that once spread to these northern regions (Kantanen et al., 2000, 2009a; Li et al., 2007). Herd books, pedigree registers and breeding associations were established in the late 19th and early 20th centuries. Early native breeds had a pivotal socioeconomic role in dairy and beef production in the northern Eurasian regions but have been almost exclusively replaced by commercial international cattle populations bred for high-input, high-output farming systems. Exceptions to this trend are Yakutian cattle in Siberia and Icelandic cattle, which continue to have high regional importance in food production (Kantanen et al., 2000, 2009a). The conservation of the genetic resources of native, typically low-profit breeds is often motivated by the fact that these breeds may possess valuable genetic variations for future animal breeding and to address the challenges that animal production will face during adaptation to future conditions, brought about by factors such as climate change (Odegård et al., 2009; Boettcher et al., 2010; Kantanen et al., 2015). In addition, breeds such as Yakutian cattle exhibit adaptation in demanding environments and may be extremely useful for enabling animal production in marginal regions (Kantanen et al., 2015).

Previous studies on the characterization of cattle genetic resources in northern Eurasian breeds have used various methods to study within-breed genetic diversity, population structure, demographic factors and interbreed relationships, e.g., autosomal and Y-chromosomal microsatellites, mitochondrial D-loop and whole-genome SNP-marker scans (Li et al., 2007; Kantanen et al., 2009b; Iso-Touru et al., 2016). These studies have indicated, for example, the genetic distinctiveness of the native northern European cattle breeds (e.g., the Finnish native breeds and Yakutian cattle) from modern commercial dairy breeds (such as the Finnish Ayrshire and Holstein breeds). In addition, a whole-genome SNP genotyping analysis detected genomic regions targeted by selection, which, for example, contain immune-related genes (Iso-Touru et al., 2016). Whole-genome sequencing (WGS)-based approaches provide additional possibilities for investigation of the genetic diversity of livestock breeds adapted to various biogeographic regions and production environments. Moreover, recent advancements in bioinformatics and statistical tools have enhanced our understanding of the demographic evolution of domestic animal species, the possible role of genomic structural variations in the adaptation of livestock breeds in the course of domestication and selection and the biological functions of these genomic variations (Gutenkunst et al., 2009; Li and Durbin, 2011; Alachiotis et al., 2012; Pavlidis et al., 2013; Wang et al., 2014b; Librado et al., 2015).

To expand our knowledge of genomic variations in northern Eurasian taurine cattle, we performed whole-genome resequencing of five animals from each of three northern native breeds, namely, Eastern Finncattle, Western Finncattle and Yakutian cattle (Figure 1). We examined the genetic diversity and population structures of the breeds and identified chromosomal regions and genes under selection pressure. We also studied the demographic history of the northern Eurasian taurine cattle by using the whole-genome sequence data.

**Figure 1.**
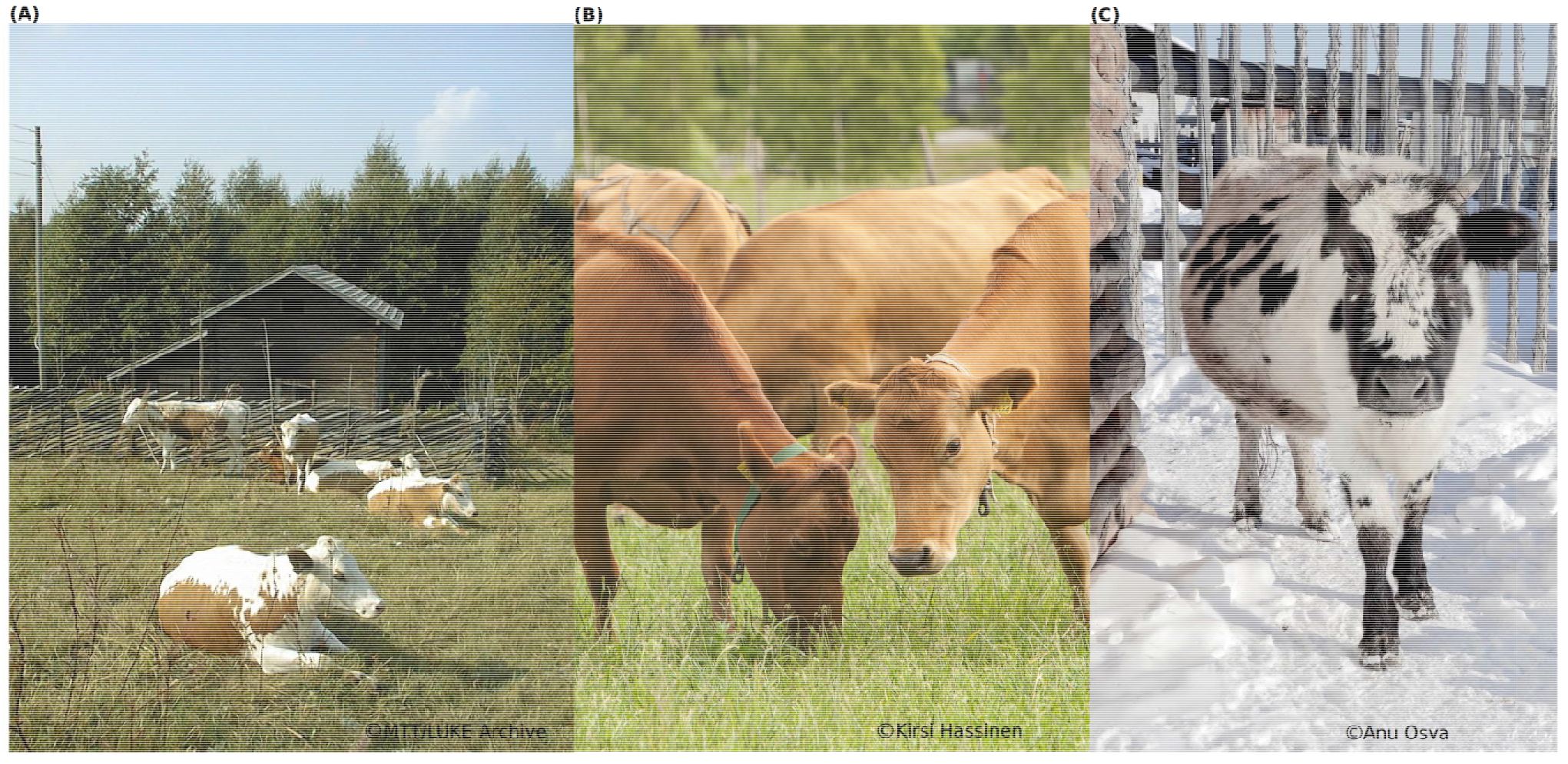
Three North Eurasian native cattle breeds are included in this study. (**A**) Eastern Finncattle are typically red-sided and polled. Cattle breeding in Finland was started with this breed, and the breed’s herd book was established in 1898. The breed was threatened with extinction in the 1970s and 1980s. The current census size is 1,600 cows, and the annual milk yield on average 4,000 Kgs. (**B**) Western Finncattle are solid light or dark brown and polled. The breed is one of the most productive native cattle breeds: the average annual milk yield is about 7,000 Kgs. (**C**) The Yakutian cattle are characterized by being purebred aboriginal native cattle from Sakha. Adult Yakutian cows weigh typically 350-400 Kgs and their height at the withers is 110-112cm on average. The animals are well adapted to Siberian harsh conditions where the temperature falls below −50°C in long winters. The average annual milk yield is 1,000 Kgs. Please do not copy, use or upload the photographs without permission of the copyright holders.

## MATERIALS AND METHODS

### Ethics statement

Blood samples of animals for DNA extraction were collected by using a protocol approved by the Animal Experiment Board of MTT Agrifood Research Finland (currently the Natural Resources Institute Finland, Luke) and the Board of Agricultural Office of Eveno-Bytantaj Region, Sakkyryr, Sakha, Russia.

### DNA sample preparation and sequencing

DNA extracted from blood samples was available for the two Finnish cattle breeds (Eastern Finncattle and Western Finncattle) and one Siberian breed (Yakutian cattle) from a previous study (Li et al., 2007). Five unrelated individuals from each breed (14 females and one Yakutian cattle bull) were examined. Genomic DNA was extracted using a standard phenol/chloroform-based protocol (Malke, 1990). For sequencing library preparation following the manufacturer’s specifications, the genomic DNA of each individual was fragmented randomly. After electrophoresis, DNA fragments of desired length were gel purified. One type of library was constructed for each sample (500 bp insert size); 15 paired-end DNA libraries were constructed for the 15 samples. Adapter ligation and DNA cluster preparation were performed, and the DNA was subjected to Illumina HiSeq 2000 sequencing using the 2 × 100 bp mode at Beijing Genomics Institute (BGI). Finally, paired-end sequence data were generated. To ensure quality, the raw data was modified by the following 2 steps: first, the contaminating adapter sequences from the reads were deleted, and then, the reads that contained more than 50% low-quality bases (quality value≤5) were removed.

### Short read alignment and mapping

For short read alignment, the bovine reference genome (UMD 3.1), including regions that were not assembled into chromosomes (Zimin et al., 2009), were downloaded from the Ensembl database release 71 (Flicek et al., 2013) and indexed using SAMtools v0.1.19 (Li et al., 2009). Paired-end 100-bp short reads from each individual sample were mapped against the bovine reference genome assembly UMD 3.1 using BWA v0.7.5a with the default parameters. After mapping, for downstream SNP and insertion-deletion (indel) detection, the SAM files that were generated from BWA were converted to the corresponding binary equivalent BAM files and sorted simultaneously using SortSam.jar in Picard tools v1.102 (http://picard.sourceforge.net/). We used Picard tools to remove PCR duplicates from the aligned reads and then used the uniquely mapped reads for variant calling.

### SNP and indel detection

We used the Genome Analysis Toolkit (GATK) v2.6-4 according to the GATK best practices pipeline (McKenna et al., 2010; DePristo et al., 2011; Van der Auwera A. et al., 2013) for downstream SNP and indel calling. We used RealignerTargetCreater to identify poorly mapped regions (nearby indels) from the alignments and realigned these regions using IndelRealigner. Next, the UnifiedGenotyper was used to call SNPs and indels with a Phred scale quality greater than 30. After SNP calling, we used VariantFiltration to discard sequencing and alignment artifacts from the SNPs with the parameters “MQ0 ≥ 4 && ((MQ0 / (1.0 * DP)) > 0.1)”, “SB ≥ −1.0, QUAL < 10”, and “QUAL < 30.0 || QD < 5.0 || HRun > 5 || SB > −0.10” and from the indels with the parameters “QD < 2.0 “, “FS > 200.0” and “ReadPosRankSum < −20.0”. All the variants that passed the above filtering criteria were used in the downstream analysis and compared to the cattle dbSNP148 (Van der Auwera A. et al., 2013) to identify novel variants.

### SNP and indel annotation and gene ontology analysis

ANNOVAR (Wang et al., 2010) was used to annotate the functions of the variants (exonic, intronic, 5’ and 3’ UTRs, splicing, intergenic) using Ensembl release 71. SNPs that were identified in the exonic regions were classified as synonymous or nonsynonymous SNPs. In recent studies, numerous phenotypes have been associated with the genes containing the highest number of nonsynonymous SNPs (nsSNPs) (Kawahara-Miki et al., 2011; Li et al., 2014). We performed gene ontology (GO) analysis for genes containing nsSNPs and indels using the GO Analysis Toolkit and Database for Agricultural Community (AgriGO) (Du et al., 2010). In this analysis, we selected genes containing >5 nsSNPs for each breed. The significantly enriched GO terms were assessed by Fisher’s exact test with the Bonferroni correction using default parameters (P-value, 0.05; at least 5 mapping entries). Out of four indel classes (frameshift, nonframeshift, stopgain and stoploss), we annotated frameshift indels in exonic regions using default parameters in ANNOVAR. Frameshift indels may change amino acid sequences and thereby affect protein function.

### Identification and annotation of selective sweeps

We investigated the signatures of selection using site frequency spectrum (SFS)-based *α* statistics in SweeD (Pavlidis et al., 2013) with default parameters, except setting the grid as the only parameter. SweeD detects the signature of selection based on the composite likelihood ratio test (CLR) using SFS-based statistics. SweeD was run separately for each chromosome by setting the grid parameter at 5-kb equidistant positions across the chromosome (size of the chromosome/5 kb). We used BEAGLE program ver.4 (Browning and Browning, 2007) to impute missing alleles and infer the haplotype phase for all individual Western Finncattle, Yakutian cattle and Eastern Finncattle simultaneously (among the Eastern Finncattle, we excluded one inbred animal; see Results). The BEAGLE program infers the haplotype information of each chromosome, which is required for *α* statistics. Following the approaches described in previous studies (Wang et al., 2014b; McManus et al., 2015), we selected the outliers falling within the top 0.5% of the CLR distribution. The cutoff value for *α* statistics was taken as the 99.5 percentile of the empirical distribution of the 5-kb equidistant positions across the genome for each chromosome. Annotation of the candidate sites that exhibited a signal of selection was performed using Ensembl BioMart (Kinsella et al., 2011) by considering a 150-kb sliding window on the outlier sites. Candidate genes exhibiting signatures of selection were subjected to GO analysis with same parameters applied in the variant annotation using AgriGO.

### Population genetics analysis

The average pairwise nucleotide diversity within a population (*π*) and the proportion of polymorphic sites (Watterson’s θ) were computed using the Bio::PopGen::Statistics package in BioPerl (v1.6.924) (Stajich et al., 2002). Principal component analysis (PCA) was conducted using smartpca in EIGENSOFT3.0 software (Patterson et al., 2006) on biallelic autosomal SNPs that were genotyped in all individuals. Significant eigenvectors were determined using Tracy Widom statistics with the twstats program implemented in the same EIGENSOFT package.

### Demographic history inference

We used the pairwise sequentially Markovian coalescent (PSMC) model (Li and Durbin, 2011) to construct the demographic history of the three breeds. For the analysis, one individual per breed with highest sequence depth was selected to explore changes in local density of heterozygous sites across the cattle genome. The following default PSMC parameters were set: −N25, −t15, −r5 and −p ‘4+25*2+4+6’. To scale the PSMC output to real time, we assumed a neutral mutation rate of 1.1 × 10-8 per generation and an average generation time of 5 years (Kumar and Subramanian, 2002; Murray et al., 2010; MacLeod et al., 2013). As the power of the PSMC approach to reconstruct recent demographic history is not reliable (Li and Durbin, 2011; MacLeod et al., 2013; Zhao et al., 2013), we reconstructed a more recent demographic history of the Finnish and Yakutian populations using the diffusion approximation for demographic inference (*∂*a*∂*i) program (dadi-1.6.3) (Gutenkunst et al., 2009). We used the intergenic sites from the identified SNPs in the 15 individuals to compute the folded SFS. We merged the results for the Eastern and Western Finncattle breeds, as these breeds exhibited similar genetic diversity measures (Figure S4). Since we had 10 Finncattle and 5 Yakutian samples, we downscaled the Finncattle sample size to be equal to that of the Yakutian cattle. We ran the *∂*a*∂*i algorithm multiple times to ensure convergence and selected the optimal parameters with the highest likelihood as the final result. As *∂*a*∂*i requires ancestral population size (Na), we calculated Na using the formula NA = θ /4μL, where θ was the observed number of segregating sites divided by the sum of the expected SFS using the best-fit parameters of our model, L was the effective sequence length, and μ was the mutation rate per generation per site. We used a mutation rate of 1.0 × 10-8 mutations per generation assuming that one generation was equal to 5 years (Kumar and Subramanian, 2002), and the effective sequence length (intergenic regions) was 10,836,904. We calculated population size and divergence time between the Finnish and Yakutian populations based on NA. Finally, using the parameters described previously, we generated the demographic model using *∂*a*∂*i as shown in Figure S5. The optimal model identified the change from the ancestral population size (NA) to the effective population size (nua) from the time Ta to the time Td. Ta was the time period when the change in NA started and Td was the time when the divergence between the Finnish and Yakutian cattle occurred. nu1F and nu2Y were the effective population sizes during the split. To calculate the statistical confidence in the estimated parameter values, we estimated the parameter uncertainties using the Hessian method (a.k.a. the Fisher information matrix).

## RESULTS

### Sequence data

A total of 521 gigabases (Gb) of paired-end DNA sequence data was obtained after removing adapter sequences and low-quality reads (Table 1, Table S1). On average, each sample had 347.4 million (M) reads, 98.45% of which were successfully mapped to the bovine reference genome UMD3.1 (Table 1, Table S1), representing 12.38-fold coverage.

**Table 1.** Summary of sequencing and short read alignment results.

|  | Eastern<br>Finn cattle | Western<br>Finn cattle | Yakutian<br>cattle | Overall<br>sample |
| --- | --- | --- | --- | --- |
| Number of individuals | 5 | 5 | 5 | 15 |
| Paired-end length (bp) | 100 | 100 | 100 | 100 |
| Average reads per individual | 352.73 M | 347.14 M | 342.33 M | 347.4 M |
| Average sequence depth per individual <sup>a</sup> | 13.21X | 13.00X | 12.82X | 13.01X |
| Average map reads per individual | 348.42 M | 340.12 M | 337.50 M | 342,02 M |
| Average unique map reads per individual | 316.89 M | 312.58 M | 309.19 M | 312.88 M |
| Average read mapping rate | 98.78% | 97.97% | 98.59% | 98.45% |
| Average coverage rate | 98.42% | 98.22% | 98.46% | 98.37% |

<sup>a</sup>Average sequence depth per individual was computed by dividing the clean reads by the reference genome size.

### Identification and annotation of variants

A total of 17.45 M SNPs were detected in the mapped reads across all 15 samples, with Yakutian cattle exhibiting the highest number of SNPs (Table 2, Figure 2a, Table S2). The average number of SNPs detected per individual within the breeds was 5.73 M, 6.03 M and 7.12 M in Eastern Finncattle, Western Finncattle and Yakutian cattle, respectively (Table S2). A total of 6.3 M (36.1%) SNPs were shared by the three breeds, and as expected, the Finnish breeds shared the highest number (n=8.06 M, 46.2%) of SNPs (Figure 1a). Moreover, we found that 1.85 M SNPs (16.83%) in Eastern Finncattle, 1.60 M (15.15%) in Western Finncattle and 3.96 M (32.33%) in Yakutian cattle were private SNPs in our data (Figure 2a). The transition-to-transversion (TS/TV) ratios were 2.20 and 2.23 in the Finncattle and Yakutian cattle, respectively (Table S2). The observed Ts/Tv ratios were consistent with those observed in previous studies in mammalian systems (Lachance et al., 2012; Choi et al., 2013, 2014), indicating the quality of our SNP data.

**Figure 2.**
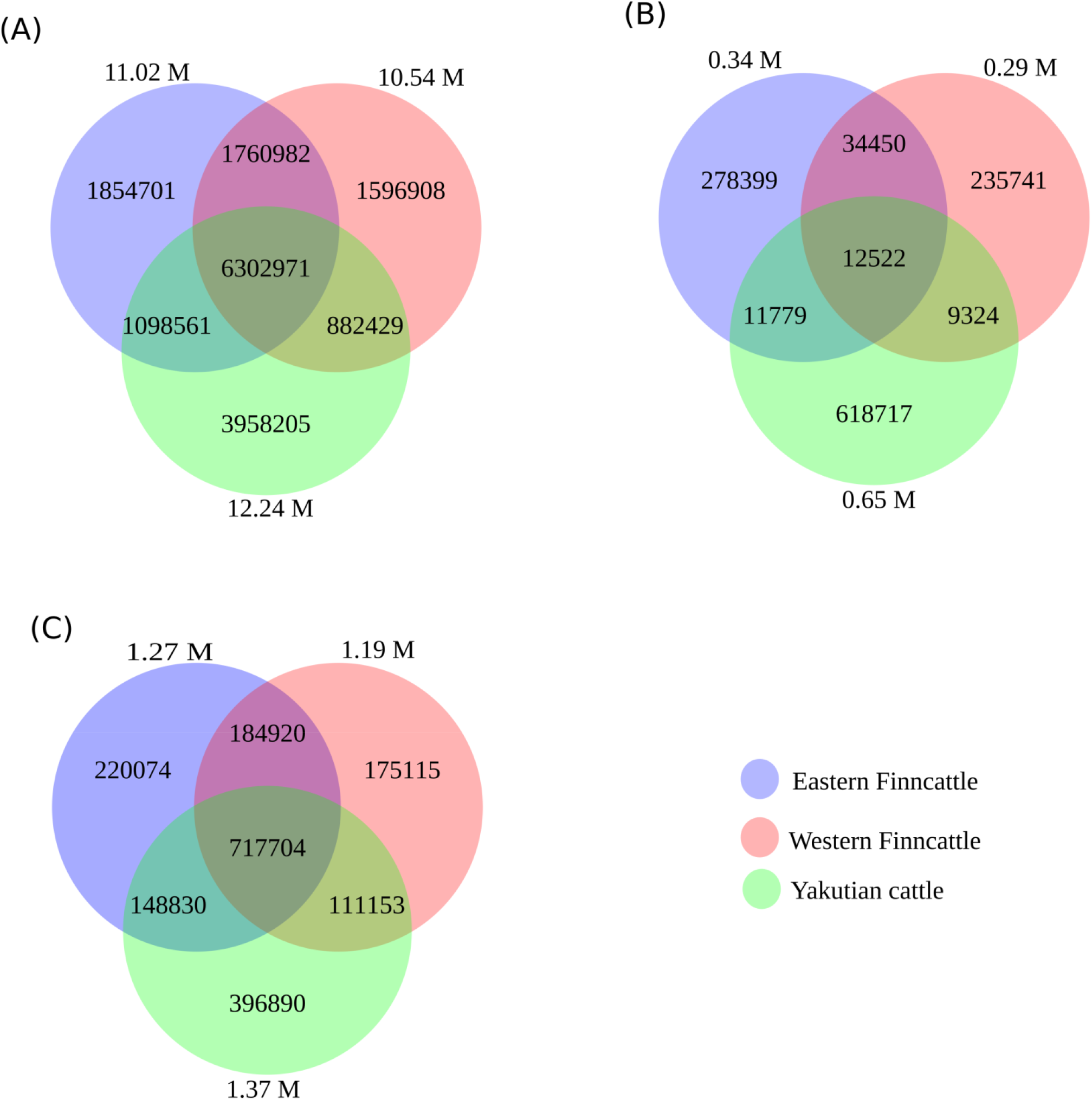
Venn diagram showing overlapping and unique SNPs/indels between the three breeds (Eastern Finncattle, Western Finncattle and Yakutian). The numbers in parentheses outside the circles are the total number of detected SNPs from each breed. The numbers in the circle components show specific SNPs for each breed or overlapping SNPs/indels between any two breeds or among three breeds. (**A**) The identified shared and specific SNPs for each breed, (**B**) the identified shared and specific novel SNPs for each breed, and (**C**) the identified shared and specific indels for each breed.

**Table 2.**
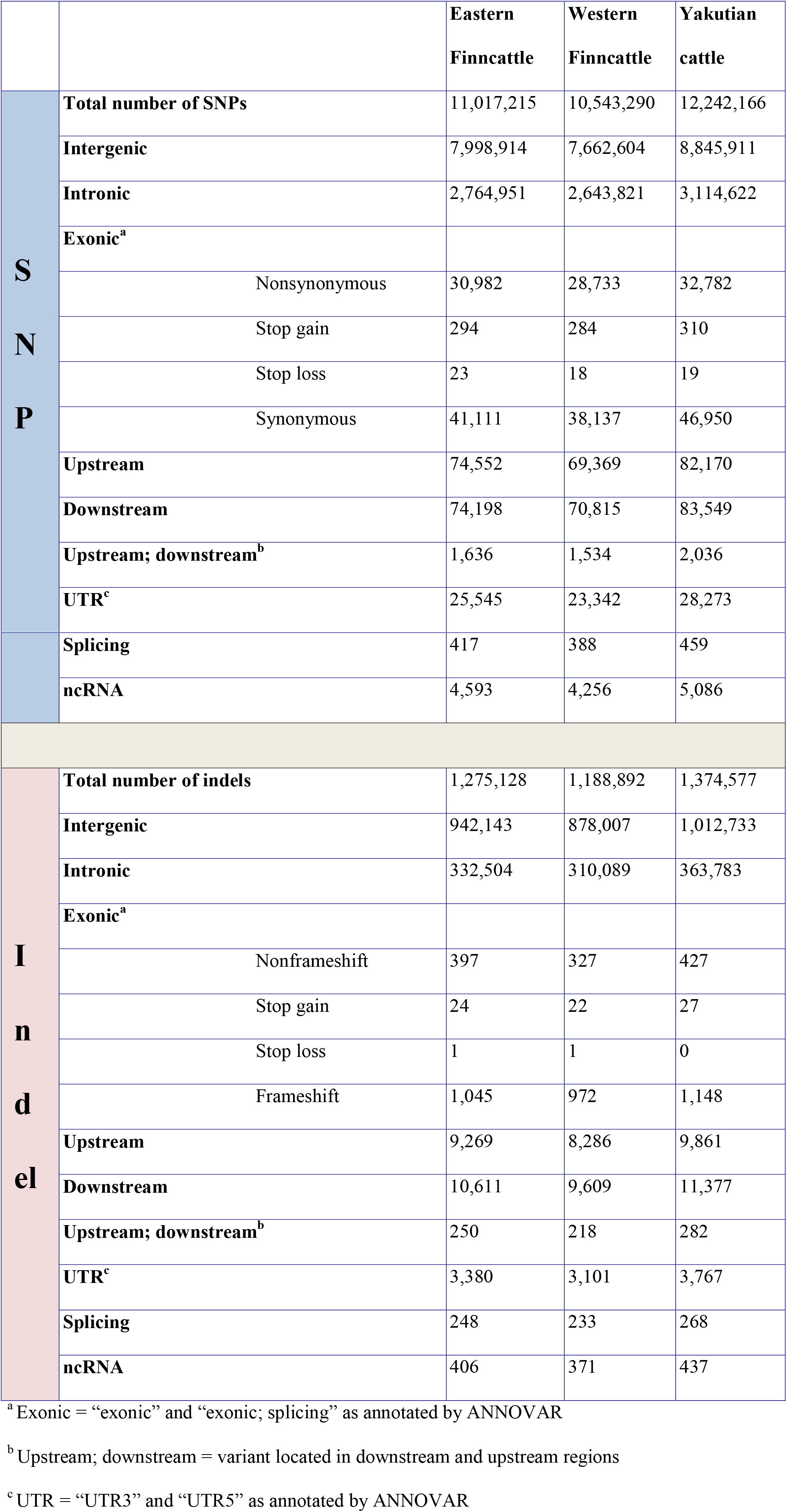
Functional annotation of the detected SNPs and indels.

Of the SNPs identified in our analysis, 1.28 M (6.9%) SNPs were found to be novel when compared to NCBI dbSNP bovine build 148. At the breed level, 3.1%, 2.8% and 5.3% of the total SNPs in the Eastern Finncattle, Western Finncattle and Yakutian cattle, respectively, were novel. Furthermore, out of the novel SNPs identified for each breed, 278,399 (82.57%), 235,741(80.72%) and 618,717 (94.85%) were breed-specific SNPs in Eastern Finncattle, Western Finncattle and Yakutian cattle (Figure 2b), respectively. A summary of the homozygous and heterozygous SNPs is given in Tables S2 and S3. One Eastern Finncattle cow (sample_3 in Table S3) exhibited exceptionally low diversity, with only 1.66 M (32.58%) heterozygous and 3.44 M (67.42%) homozygous SNPs. This animal originated from an isolated, inbred herd and represented one relict Eastern Finncattle line (herd) that passed through the breed’s demographic bottleneck (Kantanen et al., 2000). After excluding this sample, the average number of SNPs detected per Eastern Finncattle individual was 5.88 M, and the Eastern Finncattle animals exhibited 2.63 M (44.83%) homozygous and 3.24 M (55.17%) heterozygous SNPs, with a ratio of 1:1.23 (homozygous:heterozygous). Apparently, the number of homozygous SNPs in the Eastern Finncattle was higher than that in the other two breeds.

In total, we detected 2.12 M indels, 79.8% of which were found in the dbSNP build 148, with 20.2% being novel (Figure 2C, Table S2). At the breed level, 13.0%, 11.7% and 16.1% of the total indels in the Eastern Finncattle, Western Finncattle and Yakutian cattle, respectively, were novel.

In our data, on average, 0.65% of the SNPs were detected in exonic regions, 25.1% in intronic regions, 72.6% in intergenic regions, and 1.65% in UTRs and in regions upstream and downstream of genes (Table 2 and Table S4). In general, all the three breeds exhibited similar distributions of SNPs in various functional categories. A total of 76,810, 71,256 and 84,927 exonic SNPs were identified in the Eastern Finncattle, Western Finncattle and Yakutian cattle, respectively. Of the exonic SNPs in the Eastern Finncattle, Western Finncattle and Yakutian cattle, 31,299, 29,035 and 33,111, respectively, were nonsynonymous SNPs (nsSNPs) (Table 2) and were found in 10,309, 9,864 and 10,429 genes, respectively.

The functional categories of the indel mutations are presented in Table 2 and Table S4. In total, 1,045, 927 and 1,148 of the indels were frameshift indels that were associated with 808, 770 and 895 genes in Eastern Finncattle, Western Finncattle and Yakutian cattle, respectively (Supplementary Data 1, 2 and 3).

### GO analysis of the SNPs and indels

GO enrichment analysis of 1,331, 1,170 and 1,442 genes containing >5 nsSNPs (Supplementary Data 4, 5 and 6), identified 111, 113 and 95 significantly enriched GO terms in Eastern Finncattle, Western Finncattle and Yakutian cattle, respectively (Supplementary Data 7, 8 and 9). A total of 38, 43 and 38 GO terms were associated with biological processes in Eastern Finncattle, Western Finncattle and Yakutian cattle, respectively (Supplementary Data 7, 8, 9).

A detailed comparison of the biological processes associated with genes with >5 nsSNPs with the bovine Ensembl gene set (n=25,160) is shown in Figure S1. The GO enrichment analysis revealed that a majority of the significantly enriched GO terms were shared by the three cattle breeds. “Response to stimulus, GO:00050896” was associated with approximately 50% of the genes in Eastern Finncattle (n=611), Western Finncattle (n =544) and Yakutian cattle (n=629) (see Figure S1). In addition, this analysis showed that in each breed, a large number of genes were associated with immune functions, such as “Immune response, GO:0006955”, “Defense response, GO:0006952”, “Antigen processing and presentation, GO:0019882”, and “Immune system process, GO:0002376”. Among the three breeds, the Yakutian cattle had more enriched genes associated with immune functions than the two Finncattle breeds. On the other hand, in the Finncattle breeds, a large number of genes were associated with sensory perception functions, such as “Sensory perception, GO:0007600”, “Sensory perception of smell, GO:0007608” and “Detection of chemical stimulus involved in sensory perception, GO:0050907”. In Yakutian cattle, none of the GO terms associated with sensory perception were enriched. However, 55 genes associated with “Developmental growth, GO: 0048589” were enriched in only Yakutian cattle.

We further identified the top genes, namely, *TTN, PKHD1, GPR98* and *ASPM,* that had at least 40 nsSNPs in all the breeds. These genes have large sizes; *TTN* is 274 kb in size, *PKHD1* if 455 kb, *GPR98* is 188 kb and ASPM is 64 kb. Among the genes with nsSNPs, *TTN* contained the highest number of nsSNPs: 68, 63 and 87 nsSNPs in Eastern Finncattle, Western Finncattle and Yakutian cattle, respectively. The *TTN* gene is present on chromosome 2 and is associated with meat quality (Sasaki et al., 2006; Watanabe et al., 2011).

A total of 709, 675 and 772 genes associated with frameshift indels in these breeds were linked to at least one GO term (Figure S2, Supplementary Data 10, 11 and 12). The results indicated that a majority of the significantly enriched GO terms were shared by the breeds. The GO terms “Defense response, GO:0006952” and “Female pregnancy, GO:0007565” were enriched exclusively in Yakutian cattle. In total, 96 genes were enriched in “Defense response, GO:0006952”.

### Selection signatures

We identified 2,528 sites exhibiting signatures of selection in each breed, of which 58%, 61% and 53% mapped to gene regions in Eastern Finncattle, Western Finncattle and Yakutian cattle, respectively (Figure S3). Information regarding the SNPs found in selective sweep regions in each breed is shown in Table S5.

Chromosome 1 exhibited the highest (n=159) number of selection signals and chromosome 25 the lowest (n=43). Considering a 150-kb window centered on the candidate site, Western Finncattle exhibited the highest number (n=371) of candidate genes with selection signatures, followed by Eastern Finncattle (n=331), while Yakutian cattle exhibited the lowest number (n=249) (Supplementary Data 13, 14 and 15). Apparently, 36 (Eastern Finncattle), 35 (Western Finncattle) and 20 (Yakutian cattle) candidate gene IDs lacked gene descriptions (Supplementary Data 16, 17 and 18). Seven genes with greater than 5 nsSNPs in Eastern Finncattle (*CCSAP, CEP72, GBP5, LOC100297846, GBP2, LOC613867* and ENSBTAG00000045571), Western Finncattle (*CDH23, PCDHB4, PCDHB6, PCDHB7, SIRPB1, LOC783488* and ENSBTAG00000012326) and Yakutian cattle (*FER1L6, GBP5,* ENSBTAG00000015464, ENSBTAG00000025621, *GBP2,* ENSBTAG00000039016 and *LOC101902869*) exhibited the strongest signatures of selection. Of the genes with the strongest signatures of selection, one gene each from Eastern (ENSBTAG00000045571) and Western Finncattle (ENSBTAG00000012326) and three genes from Yakutian cattle (ENSBTAG00000015464, ENSBTAG00000025621, ENSBTAG00000039016) lacked gene descriptions (Table S6).

A total of 28, 67 and 13 GO terms were significantly enriched in Eastern Finncattle, Western Finncattle and Yakutian cattle, respectively (Supplementary Data 19, 20 and 21). We found only one significantly enriched GO term (“GMP binding, GO:0019002”) that was shared by the three cattle breeds. The GO terms “Homophilic cell adhesion, GO:0007156”, “Calcium-dependent cell-cell adhesion,GO:0016339” and “Multicellular organism reproduction, GO:0032504” were shared by the Finncattle breeds. Most of the significantly enriched GO terms (23, 62 and 12 in Eastern Finncattle, Western Finncattle and Yakutian cattle, respectively) were ‘breed-specific’ in our data. In addition, we examined the significantly enriched GO terms that were potentially involved in cold adaptation by assuming that in extremely cold environments, energy requirement is high and fat and lipids are the main sources of energy (Liu et al., 2014). The levels of fatty acids, lipids and phospholipids typically increase with decreasing temperatures (Purać et al., 2011). The significantly enriched GO terms associated with Western Finncattle included “Lipid localization, GO:0010876”, “Lipid digestion, GO:0044241”, “Unsaturated fatty acid biosynthetic process, GO:0006636” and “Unsaturated fatty acid metabolic process, GO:0033559 “. However, no significantly enriched GO terms associated with fatty acid and lipid metabolism and biosynthesis were identified in Eastern Finncattle and Yakutian cattle.

We examined the candidate selective sweep genes in each breed. A number of genes potentially associated with cold adaptation (Cardona et al., 2014) were present in Eastern Finncattle (*DNAJC28, HSP90B1, AGTRAP, TAF7, TRIP13, NPPA and NPPB*), Western Finncattle (*CD14, COBL, JMJD1C, KCNMA1, PLA2G4, SERPINF2, SRA1* and *TAF7*) and Yakutian cattle (*DNAJC9, SOCS3, TRPC7, SLC8A1 GLP1R, PKLR* and *TCF7L2).*

Among the selective sweep genes, there were several genes that have been previously shown to be associated with domestication-related changes, such as changes in disease resistance, neuronal and brain development, growth, meat quality, pigmentation, sensory perception and milk production (Gutiérrez et al., 2015). For example, the chromosomal regions exhibiting selective sweeps in Eastern Finncattle included genes associated with disease resistance (*IFNAR1, IFNAR2, IL10RB* and *NOD2*), neuronal and brain development (*OLIG1*), growth (*ACTA1*) and meat quality (*IGFBP5, NRAP, PC* and *S1PR1*) (Supplementary Data 13). In Western Finncattle, selective sweeps were detected in genes associated with pigmentation (*ULBP3*), sensory perception (*LOC521946, LOC783558* and *LOC783323*), meat quality (*COX5B, KAT2B* and *ITGB3*) and disease resistance (*CD96, CD14, GZMB* and *IL17A*) (Supplementary Data 14). Similarly, selective sweep-influenced genes in Yakutian cattle were associated with disease resistance (*PFKM, ADAM17* and *SIRPA*), sensory perception (*OR13C8, LOC100336881, LOC101902265, LOC512488, LOC617388, LOC783884, LOC788031* and *LOC789957*), meat quality (*ALDH1B1, CAPNS1, COX7A1, PFKM, SLC8A1, SOCS3* and *THBS3*) and milk production (*MUC1*) (Supplementary Data 15).

### Population genetics analysis

The overall genome-wide genetic diversity, as measured by Watterson’s θ and pairwise nucleotide diversity (*π*), were higher in the Yakutian cattle (0.001588 and 1.728 × 10-3, respectively) than in Eastern Finncattle (0.001445 and 1.559 × 10-3, respectively) and Western Finncattle (0.001398 and 1.512 × 10-3, respectively), and these results were inconsistent with those of previous studies based on autosomal microsatellite and SNP data sets, which showed that Finncattle were more diverse than the Yakutian cattle (Li and Kantanen, 2010).

We also applied PCA to examine the genetic relationships among the three cattle breeds. In the PCA plot, the Finncattle and Yakutian cattle were grouped in the first eigenvectors, indicating clear genetic differentiation (Figure S4). The inbred Eastern Finncattle animal grouped separately from the other Finncattle animals.

### Demographic population size history

The PSMC profiles of the contemporary Finnish and Siberian native cattle were used to construct the demographic prehistory and evolution of ancestral populations of northern Eurasian cattle. As shown in Figure 3, the temporal PSMC profiles of the three cattle genomes followed a similar pattern. The ancestral species of northern Eurasian taurine cattle, the near-eastern aurochs (*Bos primigenius*) (Kantanen et al., 2009a), experienced two population peaks starting at ~1 Mya and ~40 kya and two bottlenecks at ~250 kya and ~12 kya (Figure 3). After the first population expansion, the population size declined gradually. The second population expansion of the ancestral wild species began around ~80 kya and started to decline around ~30 kya, leading to a second bottleneck.

**Figure 3.**
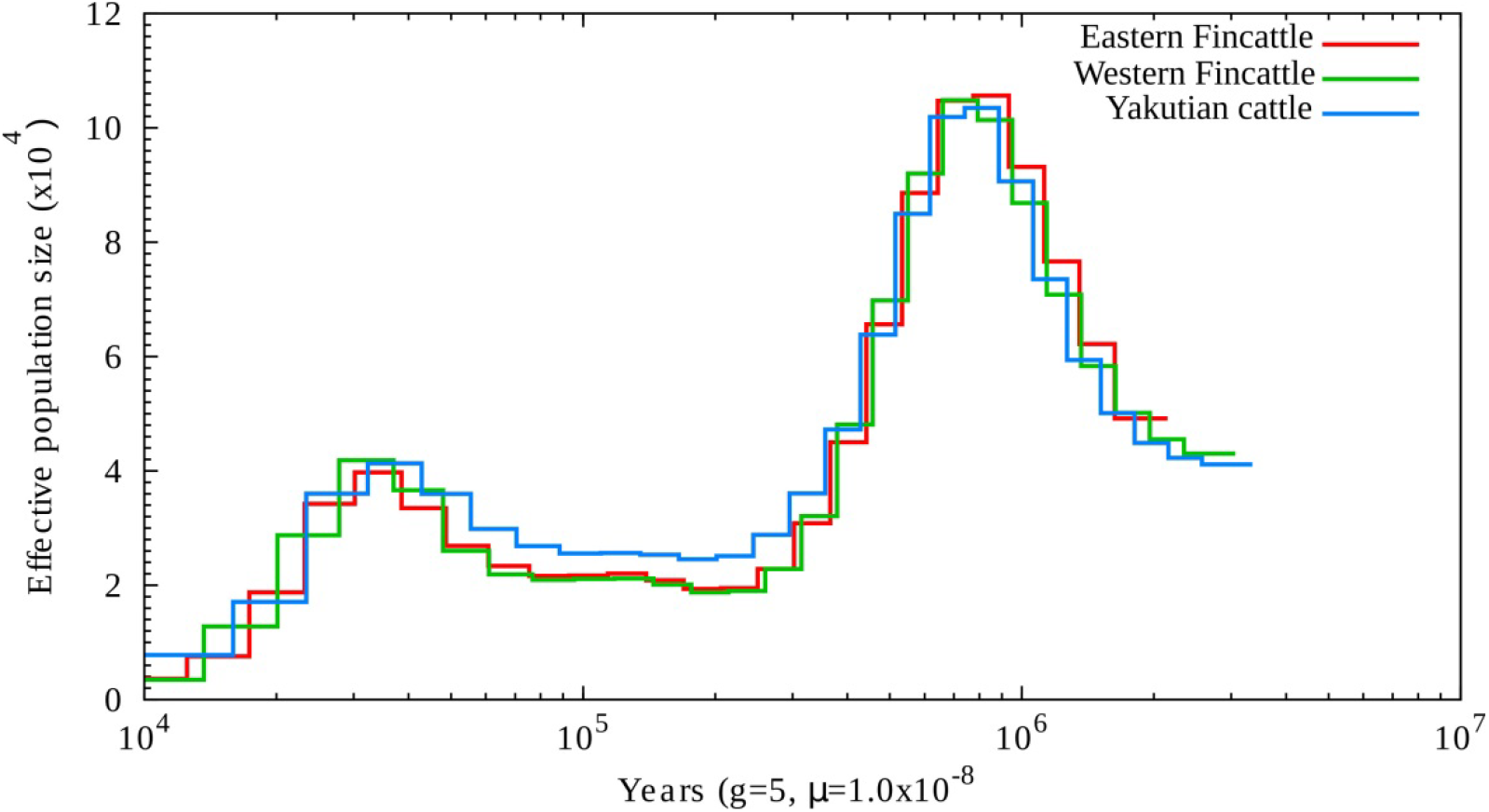
Demographic history of the northernmost cattle breed reconstructed from three cattle genomes, one from each breed, by using PSMC. The X axis shows the time in thousand years (Kyr), and the Y axis shows the effective population size.

We also used the *∂*a*∂*i program to reconstruct the recent northern European cattle demographic history (from 418 kya to the present). The parameters Ta, Td, nua, nu1F and nu2Y in the demographic model are shown and explained in Figure S5 and Table S7. Based on this model, we estimated that the reference ancestral population size (NA) was 43,116. The optimal model fit for each parameter and confidence interval (CI) are shown in Table S7 by fixing NA at 43,116 and generation time at 5 years. Our best-fit model indicated that the ancestral population underwent a size change to 51,883 (CI, 51,658-52,108) at 418 kya (95% CI, 413.96-409.47 kya) (Table S7). This result is consistent with the PSMC profile (Figure 3). In addition, our model suggested that the divergence of North European native cattle and East Siberian turano-mongolicus type of cattle occurred 8,822 years ago (CI, 8,775-8,869 years ago).

## DISCUSSION

To our knowledge, this is the first whole-genome sequence-based report on the genetic diversity of Eurasian native cattle (*B. taurus*) breeds that have adapted to the northernmost cattle farming regions, even subarctic regions. The contemporary genetic resources of the Eastern Finncattle, Western Finncattle and Yakutian cattle breeds studied are the result of a complex process of genetic and demographic events that occurred during the domestication and selection and even the evolution of the ancestral species of northern Eurasian taurine cattle, namely, the near-eastern aurochs (*B. primigenius*).

### Demographic evolution of *Bos primigenius*

As shown in Figure 3, the auroch species (*B. primigenius*) experienced two notable prehistorical population expansions, after which the population size declined gradually. The first marked decline in the effective population size (Ne) occurred during the Middle Pleistocene period starting after ~1 Mya, which may have been associated with reduction in global temperatures and even with negative actions of humans on the auroch population (Barnosky et al., 2004; Hughes et al., 2007). The second marked decline in Ne prior to domestication was obviously caused by dramatic climate changes during the last glacial maximum (Yokoyama et al., 2000). Although the sequencing depth attained in this study was not ideal for PSMC analysis (typically >20×), our observations regarding the temporal changes in the Ne of the aurochs during the Pleistocene period (Mei et al., 2018) followed the pattern observed for ancestral populations of several other domestic mammalian species, such as pig (*Sus scrofa;* (Groenen et al., 2012)), horse (*Equus caballus;* (Librado et al., 2016)) and sheep (*Ovis aries,* (Yang et al., 2016)). The *∂*a*∂*i results confirmed the past fluctuations in the prehistorical Ne of *B. primigenius* (Table S7), and the comparison of the current SNP-based estimated Ne of the present cattle breeds (~100; (Iso-Touru et al., 2016)) to the Ne of the corresponding early domesticated ancestral populations showed that there was a dramatic decline in the Ne during domestication and breed formation. In addition, our demographic analysis (Figure S5) provided new knowledge of the prehistory of northern Eurasian native cattle. As suggested by a previous study (Kantanen et al., 2009b), both the Finnish and Yakutian native cattle descended from the near-eastern aurochs domesticated 8,000-10,000 years ago. Here, our results have shown that the two northern Eurasian native cattle lineages may have already diverged in the early stage of taurine cattle domestication, more than 8,000 years ago.

### High genetic variability in the Yakutian cattle

The total number of sequence variants identified on average in Eastern Finncattle and Western Finncattle animals (e.g., 5.88 M and 6.03 M SNPs, respectively, exhibiting a minor allele frequency > 0.05) corresponded well to numbers found typically in European taurine animals. In contrast, we found that the Yakutian cattle exhibited a higher number of SNPs on average per individual (7.12 M SNPs) than the number of SNPs detected in European and Asiatic humpless cattle to date (Tsuda et al., 2013; Choi et al., 2014; Szyda et al., 2015). According to (Szyda et al., 2015) and studies cited therein, a European taurine animal may exhibit on average 2.06-6.12, 5.89-6.37, 5.85-6.40 and 5.93 M SNPs, while (Choi et al., 2014) detected 5.81M SNPs in a Korean Holstein cattle individual, a breed that originated from western Europe and North America. Typically, it may be possible to detect additional SNPs by increasing the sequencing depth (Szyda et al., 2015). In addition to the average number of SNPs per individual, total number of SNPs and number of indels, the Yakutian cattle exhibited the highest number of exonic SNPs and nsSNPs among the three northern native breeds studied. However, although the Yakutian cattle had the highest number of nsSNPs and genes with >5 nsSNPs, the functional annotation of the exonic SNPs by GO analysis indicated that the lowest number of significantly enriched GO terms was obtained for the Yakutian cattle.

Our estimates for the population-level diversity for the Eastern Finncattle, Western Finncattle and Yakutian cattle (the nucleotide diversity (*π*) values were 1.559 × 10-3, 1.512 × 10-3 and 1.728 × 103, respectively, and the proportions of polymorphic sites (θ) were 0.001445, 0.001398 and 0.001588, respectively) exceed those typically found in European taurine cattle breeds (Kim et al., 2017; Chen et al., 2018; Mei et al., 2018). We observed that Yakutian cattle such as the Asiatic taurine cattle breeds exhibit high levels of genomic diversity in terms of *π* and θ estimates. The typical nucleotide diversity values for the European taurine cattle are >1.0 × 10e-3, while those for the Asiatic taurine breeds are closer to ~2.0 × 10e-3 than to 1.0 × 10e-3 (Kim et al., 2017; Chen et al., 2018; Mei et al., 2018). We observed higher within-population diversity for the Yakutian cattle than that observed for several other taurine cattle breeds, which differs from previous estimates based on autosomal microsatellites and whole-genome SNP data (Li et al., 2007; Iso-Touru et al., 2016), where lower levels of variation were observed in Yakutian cattle, indicating that the genetic variation in Yakutian cattle has been underestimated. The set of autosomal microsatellites recommended by FAO (the Food and Agricultural Organization of the United Nations) for biodiversity analysis of cattle breeds and the design of commercial SNP BeadChips used in cattle whole-genome genotyping were derived mainly from the genetic data of western breeds, causing a bias in the diversity estimates of clearly genetically distinct cattle breeds, such as Yakutian cattle (Li et al., 2007; Iso-Touru et al., 2016).

There could have been differences in the past effective population sizes of the European and Asiatic taurine cattle, and the present elevated genomic diversity of the Asiatic taurine cattle breeds may reflect the higher “ancient” effective sizes of the ancestral populations of the Asiatic taurine breeds (Chen et al., 2018). However, the prehistory of domesticated cattle in East Asia appears to be more complex than previously thought (Zhang et al., 2013; Gao et al., 2017; Chen et al., 2018), and an additional speculative explanation for the elevated genomic diversity in the Yakutian cattle and several other Asiatic taurine cattle breeds (or their ancestral populations) could be ancient introgression with the East Asian aurochs (*B. primigenius*) that lived in the East Asian region during the arrival of near-eastern taurine cattle (Chen et al., 2018). The previous mtDNA and Y-chromosomal diversity study indicated the near-eastern origins of the ancestral population of the Yakutian cattle (Kantanen et al., 2009b). The possible hybrid origins of the Yakutian cattle ancestries may have increased the genetic variation in the ancestral population of Yakutian cattle seen even in the current population and may have played a pivotal role in the process of adaptation of the Yakutian cattle to the subarctic environment in the Sakha Republic, eastern Siberia.

The high number of SNPs and high genomic diversity found in the Yakutian cattle may be due partly to the breed’s selection history: the artificial selection by humans has not been intensive (Kantanen et al., 2009b). The Yakutian cattle breed is an aboriginal taurine population, the gene pool of which has been shaped by natural and artificial selection. However, the centuries-old “folk selection” methods and traditional knowledge for the selection of the most suitable animals for the challenging subarctic environment followed the methods used by local people rather than the breeding implemented by organizations or institutions (Kantanen et al., 2009a). When compared with the Western Finncattle and Eastern Finncattle in the present study, the Yakutian cattle exhibited distinctly low numbers of candidate genes that exhibited selection signatures (n=371, n=331 and n=249, respectively). Among these three breeds, Western Finncattle have been subjected to the most intensive artificial selection for milk production characteristics, while the production selection program of Eastern Finncattle was stopped in the 1960s, when the census population size of this native breed declined rapidly.

Currently, *in vivo* and *in vitro* conservation activities are being implemented for Eastern Finncattle (and for Western Finncattle and Yakutian cattle). In addition, although Yakutian cattle had the highest number of genes containing SNPs (also nsSNPs) among the three breeds, the GO analysis indicated that this breed had the lowest number of significantly enriched GO terms (Eastern Finncattle, 111; Western Finncattle, 113; and Yakutian cattle, 95). This difference between the native Finnish cattle and Yakutian cattle can be due to the differences in the selection histories of these breeds.

### Genomic characteristics of the northern Eurasian taurine cattle breeds

The GO enrichment analysis of genes harboring >5 nsSNPs indicated that genes related, e.g., to immunity and “response to stimulus” are overrepresented in the set of genes identified in the northern Eurasian native cattle breeds in this study. “Response to stimulus” refers to a change in the state or activity of a cell or an organism as a result of the detection of a stimulus, e.g., a change in enzyme production or gene expression (Gene Ontology Browser). This observation was consistent with previous cattle resequencing analyses (Choi et al., 2014; Stafuzza et al., 2017; Mei et al., 2018) and suggests that these genes were under positive selection during the course of cattle evolution and provided survival benefits, e.g., during environmental changes (Nielsen et al., 2007). Moreover, we observed similar GO terms in our transcriptome study that included northern Eurasian cattle breeds (Pokharel et al., 2018). Interestingly, genes related to the GO term “Sensory perception” were enriched in Eastern Finncattle and Western Finncattle but not Yakutian cattle. We performed a manual search for genes associated with “Sensory perception” genes. We found that 47 of these genes exhibited >5 nsSNPs in Eastern Finncattle and Western Finncattle, most of which were olfactory receptor genes. We determined the number of SNPs and nucleotide diversity of this set of genes and found that the Yakutian cattle exhibited less variation than the two Finnish native breeds (the number of SNPs and *π*-estimates for Eastern Finncattle, Western Finncattle and Yakutian cattle were 2,298 and 1.864 × 10e-3; 2,091 and 1.792 × 10e-3; 1,478 and 1.113 × 10e-3, respectively), which is in contrast to the number of SNPs and *π*-estimates obtained for the entire genomes of the breeds. Great variations in the number of olfactory receptor genes and structural variations in these genes among mammalian species and even individuals within species (e.g., in humans) have been interpreted as reflecting the effects of environmental factors on the genetic diversity of this multigene family and demonstrate the importance of these genes from the evolutionary point of view (Niimura, 2011; Niimura et al., 2014). Therefore, we hypothesize that the reduced genetic diversity in the evolutionarily important genes in Yakutian cattle could be associated with gradual adaptation to the challenging subarctic environment along with human movements from the southern Siberian regions to more northern environment (Librado et al., 2015). Cattle (and horses) may have been introduced to the Yakutian region after the 9th century, perhaps as late as the 13th century (Kopoteva and Partanen, 2009). Compared to European taurine cattle, this is a relatively short time period in terms of intervals between cattle generations. In our study, genes related to the GO term “Developmental growth” were enriched in only Yakutian cattle. (Stothard et al., 2011) suggested that genes associated with the GO term “Growth” may be related to the increase in the mass of intensively selected Black Angus (beef breed) and Holstein (dairy breed) cattle. However, Yakutian cattle have not been selected for increased body size as that would be less desirable characteristic in Yakutian conditions. Instead, we hypothesize that the enrichment of these growth-related genes in Yakutian cattle may be a signature of adaptation to the harsh environment. The Yakutian cattle exhibit unique morphoanatomical adaptations to the subarctic climate and are characterized by their small live weights (adult cows typically weigh 350-400 kilograms, with heights of 110-112 centimeters); deep but relatively narrow chests; and short, firm legs (Kantanen et al., 2009a). The Yakutian cattle are unique remnants of the Siberian Turano-Mongolian type of taurine cattle (Kantanen et al., 2009a) and can be distinguished from the European humpless cattle by these anatomical characteristics.

We performed genome-wide selection-mapping scans for the three northern cattle breeds and found a great majority of SNPs exhibiting selection signatures in noncoding genomic regions. This finding indicates that selection occurs specifically via the regulatory elements of genomes (see also (Librado et al., 2015)). We found that the studied breeds exhibited ‘private’ (breed-specific) selection signature patterns, indicating distinctiveness in their selection histories. We further investigated the proportions of genes exhibiting selection signatures among the breeds and found that only 5 genes from this set of genes were shared by the three breeds. Only 37 genes were shared by the two Finnish native cattle breeds, while 13 ‘selection signature’ genes were shared by Western Finncattle and Yakutian cattle and 11 by Eastern Finncattle and Yakutian cattle. In addition, the breeds did not share any of the genes exhibiting the strongest selection signatures and harboring >5 nsSNPs, and the GO term enrichment analysis of this set of genes indicated that only one GO term (“GMP binding”) was significantly enriched in all three breeds.

We identified several positively selected candidate genes underlying adaptation, appearance and production of Eastern Finncattle, Western Finncattle and Yakutian cattle. For example, in Eastern Finncattle, selection signatures were detected in *NRAP* and *IGFBP5,* both of which have been previously identified as candidate genes for muscle development and meat quality in cattle (Williams et al., 2009), and in *NOD2,* which is a candidate gene for dairy production (Ogorevc et al., 2009). In Western Finncattle, we detected selection signatures in, e.g., candidate genes for beef production, such as *COX5B* and *ITGB3* (Williams *et al.,* 2009), and dairy production, such as *CD14* (Ogorevc et al., 2009). In Yakutian cattle, several genes exhibiting selection signatures were candidate genes for muscle development and meat quality, such as *COX7A1, THBS3, PFKM,* and *SOCS3* (Williams *et al.,* 2009) but also for color pattern (*ADAM17;* (Gutiérrez-Gil et al., 2015)) and milk production traits (*MUC1;* (Ogorevc et al., 2009)). We were particularly interested in the genomic adaptation to North Eurasian environments. (Cardona et al., 2014) listed in the supplementary materials of their publication several potential candidate genes associated with biological processes and pathways hypothesized to be involved in cold adaptation in indigenous Siberian human populations in terms of response to temperature, blood pressure, basal metabolic rate, smooth muscle contraction and energy metabolism. Several of these genes also exhibited significant selection signatures in our cattle sequence data, as exemplified in the Results section of this paper. *SLC8A1* (sodium/calcium exchanger 1), influencing the oxidative stress response, is an example of the genes with significant selection signatures in Yakutian cattle, Siberian human populations (Cardona et al., 2014) and native Yakutian horses (Librado et al., 2015). This example of selection signatures and associated genes found in the Yakutian cattle and Siberian human populations (Cardona et al., 2014) indicates convergent evolution between the mammalian populations adapted to subarctic environments. Convergent evolution between mammalian species in adaptation to harsh environments has also occurred, e.g., on the Tibetan plateau, as indicated by (Wang et al., 2014a, 2015; Yang et al., 2016).

## CONCLUSIONS

We have investigated by whole-genome sequencing for the first time the genetic diversity of native cattle breeds originating from the northernmost region of cattle farming in the world. We found novel SNPs and indels and genes that have not yet been annotated. Our observations suggest that accurate reference genome assemblies are needed for genetically diverse native cattle breeds showing genetic distinctiveness, such as Yakutian cattle, in order to better understand the genetic diversity of the breeds and the effects of natural and artificial selection and adaptation. We identified a number of genes and chromosomal regions important for the adaptation and production traits of the breeds. Moreover, GO terms such as defense response, growth, sensory perception and immune response were enriched in the genes associated with selective sweeps. To improve our knowledge of the value of native breeds as genetic resources for future cattle breeding and the power of selection signature analyses, a greater number of animals of these breeds should be investigated in a wider breed diversity context.

## ABBREVIATIONS

nsSNPs: :nonsynonymous SNPs;
GO: :gene ontology;
CLR: :composite likelihood ratio;
SFS: :site frequency spectrum;
PCA: :principal component analysis;
PSMC: :pairwise sequentially Markovian coalescent;
*∂*a*∂*i: :diffusion approximation for demographic inference;
Gb: :gigabases

## DATA AVAILABILITY

The raw sequence reads (Fastq Files) for this study can be found in European Nucleotide Archive (ENA) under the accession number PRJEB28185 (please see Table S1 for sample specific accessions).

## AUTHOR CONTRIBUTIONS

JK designed the study, and revised the manuscript. MW performed the bioinformatics and statistical analyses and drafted the manuscript. JK, RP, IA and ZI collected the samples. RP, KP, IA, MY and ZI participated in the experimental design and paper revision. All authors read and approved the final manuscript.

## ACKNOWLEDGMENTS

We thank Tuula-Marjatta Hamama for DNA extraction and BGI for sequencing the samples. The authors wish to acknowledge the CSC-IT Center for Science, Finland, for computational resources. The owners of the animals included in the study are acknowledged for providing samples for this study.

## REFERENCES

Alachiotis, N., Stamatakis, A., and Pavlidis, P. (2012). OmegaPlus: a scalable tool for rapid detection of selective sweeps in whole-genome datasets. Bioinformatics 28, 2274–2275. doi:10.1093/bioinformatics/bts419.

Barnosky, A. D., Koch, P. L., Feranec, R. S., Wing, S. L., and Shabel, A. B. (2004). Assessing the Causes of Late Pleistocene Extinctions on the Continents. Science (80-.). 306, 70. Available at: http://science.sciencemag.org/content/306/5693/70.abstract.

Bläuer, A., and Kantanen, J. (2013). Transition from hunting to animal husbandry in Southern, Western and Eastern Finland: new dated osteological evidence. J. Archaeol. Sci. 40, 1646–1666. doi:http://dx.doi.org/10.1016/j.jas.2012.10.033.

Boettcher, P. J., Tixier-Boichard, M., Toro, M. A., Simianer, H., Eding, H., Gandini, G., et al. (2010). Objectives, criteria and methods for using molecular genetic data in priority setting for conservation of animal genetic resources. Anim. Genet. 41, 64–77. doi:10.1111/j.1365-2052.2010.02050.x.

Browning, S. R., and Browning, B. L. (2007). Rapid and Accurate Haplotype Phasing and Missing-Data Inference for Whole-Genome Association Studies By Use of Localized Haplotype Clustering. Am. J. Hum. Genet. 81, 1084–1097. Available at: http://www.ncbi.nlm.nih.gov/pmc/articles/PMC2265661/.

Cardona, A., Pagani, L., Antao, T., Lawson, D. J., Eichstaedt, C. A., Yngvadottir, B., et al. (2014). Genome-Wide Analysis of Cold Adaptation in Indigenous Siberian Populations. PLoS One 9, e98076. doi:10.1371/journal.pone.0098076.

Chen, N., Cai, Y., Chen, Q., Li, R., Wang, K., Huang, Y., et al. (2018). Whole-genome resequencing reveals world-wide ancestry and adaptive introgression events of domesticated cattle in East Asia. Nat. Commun. 9, 2337. doi:10.1038/s41467-018-04737-0.

Choi, J.-W., Liao, X., Park, S., Jeon, H.-J., Chung, W.-H., Stothard, P., et al. (2013). Massively Parallel Sequencing of Chikso (Korean Brindle Cattle) to Discover Genome-Wide SNPs and InDels. Mol. Cells 36, 203–211. doi:10.1007/s10059-013-2347-0.

Choi, J.-W., Liao, X., Stothard, P., Chung, W.-H., Jeon, H.-J., Miller, S. P., et al. (2014). Whole-Genome Analyses of Korean Native and Holstein Cattle Breeds by Massively Parallel Sequencing. PLoS One 9, e101127. Available at: http://dx.doi.org/10.1371%2Fjournal.pone.0101127.

Cramp, L. J. E., Jones, J., Sheridan, A., Smyth, J., Whelton, H., Mulville, J., et al. (2014). Immediate replacement of fishing with dairying by the earliest farmers of the northeast Atlantic archipelagos. Proc. R. Soc. B Biol. Sci. 281, 20132372. doi:10.1098/rspb.2013.2372.

DePristo, M. A., Banks, E., Poplin, R. E., Garimella, K. V, Maguire, J. R., Hartl, C., et al. (2011). A framework for variation discovery and genotyping using next-generation DNA sequencing data. Nat. Genet. 43, 491–498. doi:10.1038/ng.806.

Du, Z., Zhou, X., Ling, Y., Zhang, Z., and Su, Z. (2010). agriGO: a GO analysis toolkit for the agricultural community. Nucleic Acids Res. 38, W64–W70. doi:10.1093/nar/gkq310.

Egorov, E. G., Nikiforov, M. M., and Danilov, Y. G. (2015). Unique experience and achievements of sakha people in development of agriculture in the north. 9.

Felius, M. (1995). Cattle Breeds, an Encyclopedia; Misset Uitgeverij: Doetinchem, The Netherlands.

Flicek, P., Ahmed, I., Amode, M. R., Barrell, D., Beal, K., Brent, S., et al. (2013). Ensembl 2013. Nucleic Acids Res. 41, D48–D55. doi:10.1093/nar/gks1236.

Gao, Y., Gautier, M., Ding, X., Zhang, H., Wang, Y., Wang, X., et al. (2017). Species composition and environmental adaptation of indigenous Chinese cattle. Sci. Rep. 7, 16196. doi:10.1038/s41598-017-16438-7.

Groenen, M. A. M., Archibald, A. L., Uenishi, H., Tuggle, C. K., Takeuchi, Y., Rothschild, M. F., et al. (2012). Analyses of pig genomes provide insight into porcine demography and evolution. Nature 491, 393–398. doi:10.1038/nature11622.

Gutenkunst, R. N., Hernandez, R. D., Williamson, S. H., and Bustamante, C. D. (2009). Inferring the Joint Demographic History of Multiple Populations from Multidimensional SNP Frequency Data. PLoS Genet 5, e1000695. Available at: http://dx.doi.org/10.1371%2Fjournal.pgen.1000695.

Gutiérrez-Gil, B., Arranz, J. J., and Wiener, P. (2015). An interpretive review of selective sweep studies in Bos taurus cattle populations: identification of unique and shared selection signals across breeds. Front. Genet. 6, 167. Available at: https://www.frontiersin.org/article/10.3389/fgene.2015.00167.

Hughes, P. D., Woodward, J. C., and Gibbard, P. L. (2007). Middle Pleistocene cold stage climates in the Mediterranean: New evidence from the glacial record. Earth Planet. Sci. Lett. 253, 50–56. doi: https://doi.org/10.1016/j.epsl.2006.10.019.

Iso-Touru, T., Tapio, M., Vilkki, J., Kiseleva, T., Ammosov, I., Ivanova, Z., et al. (2016). Genetic diversity and genomic signatures of selection among cattle breeds from Siberia, eastern and northern Europe. Anim. Genet. 47, 647–657. doi:10.1111/age.12473.

Kantanen, J., Ammosov, I., Li, M., Osva, A., and Popov, R. (2009a). “A cow of the permafrost,” in Sakha Ynaga□: cattle of the Yakuts (Helsinki: Suomalaisen tiedeakatemian toimituksia), 19–44.

Kantanen, J., Edwards, C. J., Bradley, D. G., Viinalass, H., Thessler, S., Ivanova, Z., et al. (2009b). Maternal and paternal genealogy of Eurasian taurine cattle (Bos taurus). Heredity (Edinb). 103, 404–415. Available at: http://www.nature.com/hdy/journal/v103/n5/suppinfo/hdy200968s1.html.

Kantanen, J., Låvendahl, P., Strandberg, E., Eythorsdottir, E., Li, M.-H., Kettunen-PrÃ¦bel, A., et al. (2015). Utilization of farm animal genetic resources in a changing agro-ecological environment in the Nordic countries. Front. Genet. 6, 52. doi:10.3389/fgene.2015.00052.

Kantanen, J., Olsaker, I., Holm, L.-E., Lien, S., Vilkki, J., Brusgaard, K., et al. (2000). Genetic diversity and population structure of 20 north European cattle breeds. J. Hered. 91, 446–457. doi:10.1093/jhered/91.6.446.

Kawahara-Miki, R., Tsuda, K., Shiwa, Y., Arai-Kichise, Y., Matsumoto, T., Kanesaki, Y., et al. (2011). Whole-genome resequencing shows numerous genes with nonsynonymous SNPs in the Japanese native cattle Kuchinoshima-Ushi. BMC Genomics 12, 103. Available at: http://www.biomedcentral.com/1471-2164/12/103.

Kim, J., Hanotte, O., Mwai, O. A., Dessie, T., Bashir, S., Diallo, B., et al. (2017). The genome landscape of indigenous African cattle. Genome Biol. 18, 34. doi:10.1186/s13059-017-1153-y.

Kinsella, R. J., KÃ¤hÃ¤ri, A., Haider, S., Zamora, J., Proctor, G., Spudich, G., et al. (2011). Ensembl BioMarts: a hub for data retrieval across taxonomic space. Database 2011, bar030–bar030. Available at: http://dx.doi.org/10.1093/database/bar030.

Kopoteva, I., and Partanen, U. (2009). “Sakha Ynaga□: Cattle of the Yakuts.,” in A historical excursion to Northern Sakha., eds. J. Granberg, K. Soini, and J. Kantanen (Helsinki: Suomalainen Tiedeakatemia), 75–116.

Kumar, S., and Subramanian, S. (2002). Mutation rates in mammalian genomes. Proc. Natl. Acad. Sci. 99, 803–808. doi:10.1073/pnas.022629899.

Lachance, J., Vernot, B., Elbers, C. C., Ferwerda, B., Froment, A., Bodo, J.-M., et al. (2012). Evolutionary history and adaptation from high-coverage whole-genome sequences of diverse African hunter-gatherers. Cell 150, 457–469. doi:10.1016/j.cell.2012.07.009.

Li, H., and Durbin, R. (2011). Inference of human population history from individual whole-genome sequences. Nature 475, 493–496. Available at: http://www.nature.com/nature/journal/v475/n7357/abs/nature10231.html#supplementary-information.

Li, H., Handsaker, B., Wysoker, A., Fennell, T., Ruan, J., Homer, N., et al. (2009). The Sequence Alignment/Map format and SAMtools. Bioinformatics 25, 2078–2079. doi:10.1093/bioinformatics/btp352.

Li, M.-H., and Kantanen, J. (2010). Genetic structure of Eurasian cattle (*Bos taurus*) based on microsatellites: clarification for their breed classification. Anim. Genet. 41, 150–158. doi:10.1111/j.1365-2052.2009.01980.x.

Li, M., Tapio, I., Vilkki, J., Ivanova, Z., Kiselyova, T., Marzanov, N., et al. (2007). The genetic structure of cattle populations (Bos taurus) in northern Eurasia and the neighbouring Near Eastern regions: implications for breeding strategies and conservation. Mol. Ecol. 16, 3839–3853. doi:10.1111/j.1365-294X.2007.03437.x.

Li, M., Tian, S., Yeung, C. K. L., Meng, X., Tang, Q., Niu, L., et al. (2014). Whole-genome sequencing of Berkshire (European native pig) provides insights into its origin and domestication. Sci.Rep. 4. Available at: http://10.0.4.14/srep04678.

Librado, P., Der Sarkissian, C., Ermini, L., Schubert, M., Jónsson, H., Albrechtsen, A., et al. (2015). Tracking the origins of Yakutian horses and the genetic basis for their fast adaptation to subarctic environments. Proc. Natl. Acad. Sci. 112, E6889–E6897. doi:10.1073/pnas.1513696112.

Librado, P., Fages, A., Gaunitz, C., Leonardi, M., Wagner, S., Khan, N., et al. (2016). The Evolutionary Origin and Genetic Makeup of Domestic Horses. Genetics 204, 423–434. doi:10.1534/genetics.116.194860.

Liu, S., Lorenzen, E. D., Fumagalli, M., Li, B., Harris, K., Xiong, Z., et al. (2014). Population Genomics Reveal Recent Speciation and Rapid Evolutionary Adaptation in Polar Bears. Cell 157, 785–794. doi:10.1016/j.cell.2014.03.054.

MacLeod, I. M., Larkin, D. M., Lewin, H. A., Hayes, B. J., and Goddard, M. E. (2013). Inferring Demography from Runs of Homozygosity in Whole-Genome Sequence, with Correction for Sequence Errors. Mol. Biol. Evol. 30, 2209–2223. doi:10.1093/molbev/mst125.

Malke, H. (1990). J. Sambrock, E. F. Fritsch and T. Maniatis, Molecular Cloning, A Laboratory Manual (Second Edition), Volumes 1, 2 and 3. 1625 S., zahlreiche Abb. und Tab. Cold Spring Harbor 1989. Cold Spring Harbor Laboratory Press. $ 115.00. ISBN: 0-87969-309-6. J. Basic Microbiol. 30, 623. doi:10.1002/jobm.3620300824.

McKenna, A., Hanna, M., Banks, E., Sivachenko, A., Cibulskis, K., Kernytsky, A., et al. (2010). The Genome Analysis Toolkit: A MapReduce framework for analyzing next-generation DNA sequencing data. Genome Res. 20, 1297–1303. doi:10.1101/gr.107524.110.

McManus, K. F., Kelley, J. L., Song, S., Veeramah, K. R., Woerner, A. E., Stevison, L. S., et al. (2015). Inference of Gorilla Demographic and Selective History from Whole-Genome Sequence Data. Mol. Biol. Evol. 32, 600–612. doi:10.1093/molbev/msu394.

Mei, C., Wang, H., Liao, Q., Wang, L., Cheng, G., Wang, H., et al. (2018). Genetic Architecture and Selection of Chinese Cattle Revealed by Whole Genome Resequencing. Mol. Biol. Evol. 35, 688–699. doi:10.1093/molbev/msx322.

Murray, C., Huerta-Sanchez, E., Casey, F., and Bradley, D. G. (2010). Cattle demographic history modelled from autosomal sequence variation. Philos. Trans. R. Soc. London B Biol. Sci. 365, 2531–2539. Available at: http://rstb.royalsocietypublishing.org/content/365/1552/2531.abstract.

Nielsen, R., Hellmann, I., Hubisz, M., Bustamante, C., and Clark, A. G. (2007). Recent and ongoing selection in the human genome. Nat. Rev. Genet. 8, 857–868. doi:10.1038/nrg2187.

Niimura, Y. (2011). Olfactory Receptor Multigene Family in Vertebrates: From the Viewpoint of Evolutionary Genomics. Curr. Genomics 13, 103–114. doi:10.2174/138920212799860706.

Niimura, Y., Matsui, A., and Touhara, K. (2014). Extreme expansion of the olfactory receptor gene repertoire in African elephants and evolutionary dynamics of orthologous gene groups in 13 placental mammals. Genome Res. 24, 1485–1496. doi:10.1101/gr.169532.113.

Odegård, J., Yazdi, M. H., Sonesson, A. K., and Meuwissen, T. H. E. (2009). Incorporating desirable genetic characteristics from an inferior into a superior population using genomic selection. Genetics 181, 737–45. doi:10.1534/genetics.108.098160.

Ogorevc, J., Kunej, T., Razpet, A., and Dovc, P. (2009). Database of cattle candidate genes and genetic markers for milk production and mastitis. Anim. Genet. 40, 832–851. doi:10.1111/j.1365-2052.2009.01921.x.

Patterson, N., Price, A. L., and Reich, D. (2006). Population Structure and Eigenanalysis. PLoS Genet 2, e190. Available at: http://dx.plos.org/10.1371%2Fjournal.pgen.0020190.

Pavlidis, P., Živkovic, D., Stamatakis, A., and Alachiotis, N. (2013). SweeD: Likelihood-Based Detection of Selective Sweeps in Thousands of Genomes. Mol. Biol. Evol. 30, 2224–2234. doi:10.1093/molbev/mst112.

Pokharel, K., Weldenegodguad, M., Popov, R., Honkatukia, M., Huuki, H., Lindeberg, H., et al. (2018). Whole blood transcriptome analysis reveals footprints of cattle adaptation to sub-arctic conditions. bioRxiv, 379925. doi:10.1101/379925.

Purać, J., Pond, D. W., Grubor-Lajšić, G., Kojić, D., Blagojević, D. P., Worland, M. R., et al. (2011). Cold hardening induces transfer of fatty acids between polar and nonpolar lipid pools in the Arctic collembollan Megaphorura arctica. Physiol. Entomol. 36, 135–140. doi:10.1111/j.1365-3032.2010.00772.x.

Sasaki, Y., Nagai, K., Nagata, Y., Doronbekov, K., Nishimura, S., Yoshioka, S., et al. (2006). Exploration of genes showing intramuscular fat deposition-associated expression changes in musculus longissimus muscle. Anim. Genet. 37, 40–46. doi:10.1111/j.1365-2052.2005.01380.x.

Stafuzza, N. B., Zerlotini, A., Lobo, F. P., Yamagishi, M. E. B., Chud, T. C. S., Caetano, A. R., et al. (2017). Single nucleotide variants and InDels identified from whole-genome re-sequencing of Guzerat, Gyr, Girolando and Holstein cattle breeds. PLoS One 12, e0173954. Available at: https://doi.org/10.1371/journal.pone.0173954.

Stajich, J. E., Block, D., Boulez, K., Brenner, S. E., Chervitz, S. A., Dagdigian, C., et al. (2002). The Bioperl Toolkit: Perl Modules for the Life Sciences. Genome Res. 12, 1611–1618. doi:10.1101/gr.361602.

Stothard, P., Choi, J.-W., Basu, U., Sumner-Thomson, J. M., Meng, Y., Liao, X., et al. (2011). Whole genome resequencing of black Angus and Holstein cattle for SNP and CNV discovery. BMC Genomics 12, 559. doi:10.1186/1471-2164-12-559.

Szyda, J., FrÄ…szczak, M., Mielczarek, M., Giannico, R., Minozzi, G., Nicolazzi, E. L., et al. (2015). The assessment of inter-individual variation of whole-genome DNA sequence in 32 cows. Mamm. Genome 26, 658–665. doi:10.1007/s00335-015-9606-7.

Tsuda, K., Kawahara-Miki, R., Sano, S., Imai, M., Noguchi, T., Inayoshi, Y., et al. (2013). Abundant sequence divergence in the native Japanese cattle Mishima-Ushi (Bos taurus) detected using whole-genome sequencing. Genomics 102, 372–378. doi:10.1016/j.ygeno.2013.08.002.

Van der Auwera A., G., Carneiro, M. O., Hartl, C., Poplin, R., del Angel, G., Levy-Moonshine, A., et al. (2013). From FastQ data to high confidence variant calls: the Genome Analysis Toolkit best practices pipeline. Curr. Protoc. Bioinforma. / Ed. board, Andreas D. Baxevanis…[et al.] 11, 11.10.1–11.10.33. doi:10.1002/0471250953.bi1110s43.

Wang, G.-D., Fan, R.-X., Zhai, W., Liu, F., Wang, L., Zhong, L., et al. (2014a). Genetic Convergence in the Adaptation of Dogs and Humans to the High-Altitude Environment of the Tibetan Plateau. Genome Biol. Evol. 6, 2122–2128. Available at: http://dx.doi.org/10.1093/gbe/evu162.

Wang, K., Li, M., and Hakonarson, H. (2010). ANNOVAR: functional annotation of genetic variants from high-throughput sequencing data. Nucleic Acids Res. 38, e164–e164. doi:10.1093/nar/gkq603.

Wang, M.-S., Li, Y., Peng, M.-S., Zhong, L., Wang, Z.-J., Li, Q.-Y., et al. (2015). Genomic Analyses Reveal Potential Independent Adaptation to High Altitude in Tibetan Chickens. Mol. Biol. Evol. 32, 1880–1889. Available at: http://dx.doi.org/10.1093/molbev/msv071.

Wang, M., Yu, Y., Haberer, G., Marri, P. R., Fan, C., Goicoechea, J. L., et al. (2014b). The genome sequence of African rice (Oryza glaberrima) and evidence for independent domestication. Nat. Genet. 46, 982–988. Available at: http://10.0.4.14/ng.3044.

Watanabe, N., Satoh, Y., Fujita, T., Ohta, T., Kose, H., Muramatsu, Y., et al. (2011). Distribution of allele frequencies at TTN g.231054C > T, RPL27A g.3109537C > T and AKIRIN2 c.*188G > A between Japanese Black and four other cattle breeds with differing historical selection for marbling. BMC Res. Notes 4, 10. Available at: http://www.biomedcentral.com/1756-0500/4/10.

Williams, J. L., Dunner, S., Valentini, A., Mazza, R., Amarger, V., Checa, M. L., et al. (2009). Discovery, characterization and validation of single nucleotide polymorphisms within 206 bovine genes that may be considered as candidate genes for beef production and quality. Anim. Genet. 40, 486–491. doi:10.1111/j.1365-2052.2009.01874.x.

Yang, J., Li, W.-R., Lv, F.-H., He, S.-G., Tian, S.-L., Peng, W.-F., et al. (2016). Whole-Genome Sequencing of Native Sheep Provides Insights into Rapid Adaptations to Extreme Environments. Mol. Biol. Evol. 33, 2576–2592. doi:10.1093/molbev/msw129.

Yokoyama, Y., Lambeck, K., De Deckker, P., Johnston, P., and Fifield, L. K. (2000). Timing of the Last Glacial Maximum from observed sea-level minima. Nature 406, 713. Available at: http://10.0.4.14/35021035.

Zhang, H., Paijmans, J. L. A., Chang, F., Wu, X., Chen, G., Lei, C., et al. (2013). Morphological and genetic evidence for early Holocene cattle management in northeastern China. Nat. Commun. 4, 2755. Available at: http://10.0.4.14/ncomms3755.

Zhao, S., Zheng, P., Dong, S., Zhan, X., Wu, Q., Guo, X., et al. (2013). Whole-genome sequencing of giant pandas provides insights into demographic history and local adaptation. Nat. Genet. 45, 67–71. Available at: http://www.nature.com/ng/journal/v45/n1/abs/ng.2494.html#supplementary-information.

Zimin, A., Delcher, A., Florea, L., Kelley, D., Schatz, M., Puiu, D., et al. (2009). A whole-genome assembly of the domestic cow, Bos taurus. Genome Biol. 10, R42. Available at: http://genomebiology.com/2009/10/4/R42.

